# Identification of Genes Regulating Cell Death in *Staphylococcus aureus*

**DOI:** 10.1101/569053

**Authors:** Rebecca Yee, Jie Feng, Jiou Wang, Jiazhen Chen, Ying Zhang

## Abstract

*Staphylococcus aureus* is an opportunistic pathogen that causes acute and chronic infections. Due to *S. aureus*’ s highly resistant and persistent nature, it is paramount to identify better drug targets in order to eradicate *S. aureus* infections. Despite the efforts in understanding bacterial cell death, the genes and pathways of *S. aureus* cell death remain elusive. Here, we performed a genome-wide screen using a transposon mutant library to study the genetic mechanisms involved in *S. aureus* cell death. Using a precisely controlled heat-ramp and acetic acid exposure assays, mutations in 27 core genes (*hsdR1, hslO, nsaS, sspA, folD, mfd, vraF, kdpB*, USA300HOU_2684, 0868, 0369, 0420, 1154, 0142, 0930, 2590, 0997, 2559, 0044, 2004, 1209, 0152, 2455, 0154, 2386, 0232, 0350 involved in transporters, transcription, metabolism, peptidases, kinases, transferases, SOS response, nucleic acid and protein synthesis) caused the bacteria to be more death-resistant. In addition, we identified mutations in core 10 genes (*capA, gltT, mnhG1*,USA300HOU_1780, 2496, 0200, 2029, 0336, 0329, 2386, involved in transporters, metabolism, transcription, cell wall synthesis) from heat-ramp and acetic acid that caused the bacteria to be more death-sensitive or with defect in persistence. Interestingly, death-resistant mutants were more virulent than the parental strain USA300 and caused increased mortality in a *Caenorhabditis elegans* infection model. Conversely, death-sensitive mutants were less persistent and formed less persister cells upon exposure to different classes of antibiotics. These findings provide new insights into the mechanisms of *S. aureus* cell death and offer new therapeutic targets for developing more effective treatments caused by *S. aureus*.

## Introduction

*S. aureus* is a major bacterial pathogen that colonizes on the skin of over one-third of the human population and can cause acute infections such as bacteremia, pneumonia, meningitis and persistent infections such as osteomyelitis, endocarditis, and biofilm infections such as on prosthetic implants [1]. Due to emerging resistance and high risk of nosocomial infections and community-acquired infections, *S. aureus* infections are a major public health concern. To develop better treatments for *S. aureus* and other bacterial infections, understanding how bacterial cells die is crucial.

By definition, bacteria treated with bactericidal antibiotics, such as the β-lactams, quinolones, and aminoglycosides, are killed. Much is known about how the drug makes contact with its target in the bacterial cell. β-lactam antibiotics bind to penicillin binding protein (PBP) disrupting proper cell wall synthesis; quinolones bind to toposismerase or gyrase blocking DNA replication, and aminoglycosides bind to ribosomal proteins resulting in mistranslated proteins [2–4]. Despite having different drug-target interactions, the downstream effects of drug lethality of cidal antibiotics are similar. It has been proposed that bacteria treated with cidal antibiotics have pathways such as SOS response, TCA cycle, ROS formation damaging Fe-S cluster regardless of the drug target [5, 6], however, the ROS theory of cidal antibiotic lethality has been challenged as killing of bacteria can still occur in apparently anaerobic conditions that do not produce ROS [7–9].

Mechanisms pertaining to bacterial cell death were mainly characterized in toxin-antitoxin systems. One of the better characterized Toxin-Antitoxin modules in the context of bacterial cell death is the MazEF module found in *E. coli* and other species such as *Listeria, Enterococcus, Neisseria, Streptococcus* and *Mycobacterium* [10–13]. Upon exposure to stresses such as nutrient depletion, DNA damage, temperature, antibiotics, and oxidative radicals, the MazF antitoxin is degraded and hence, the MazE toxin can degrade cellular mRNA causing cellular shutdown [10, 12]. In particular to *S. aureus*, the CidA and LrgA proteins, which are holin-like molecules with analogous functions to apoptotic regulators of the BCL-2 family in eukaryotes, were proposed to play a role in death and lysis of *S. aureus* [14, 15]. However, the specific process as to how CidA and LrgA regulate cell death is poorly defined.

Although cell death is an important process of a living organism, little is known about the mechanisms. High-throughput screens have been developed to study the cell death mechanism of unicellular eukaryotic organisms such as *S. cerevisiae* upon stress signals from high temperature and acetic acid [16–19], both of which also induce death in bacteria. Here, we performed a high-throughput genetic screen using a transposon mutant library of USA300 to identify genes involved in cell death in *S. aureus* [20]. Under multiple death stimuli, we identified 27 genes whose mutations caused the bacteria to be more death-resistant, while mutations in 10 genes caused the bacteria to be more death-sensitive.

## Results

### Identification of genes important for cell death resistance

To better understand the mechanisms of cell death, we performed a genetic screen using the Nebraska Transposon Mutant Library (NTML) which contains mutations in all the non-essential genes of *S. aureus* USA300, the most common circulating community-acquired MRSA strain in the United States [20]. To design our assay, we utilized the heat-ramp assay that has been used to study cell death programs in yeast [17]. To determine viability, we employed both the traditional agar replica plating for visualization of viable growth on solid media but also stained cells with SYBR Green I/PI, a viability stain that can detect both live and dead cells [21, 22]. Using a *cidA* mutant [15] which has been shown to be death-resistant as a control and the parental strain of USA300 as a death sensitive control, we optimized the condition of our heat-ramp experiment to show the biggest difference between both the death-resistant and death-sensitive phenotypes based on agar plating and the live/dead ratio from viability staining with SYBR Green I/PI.

For identification of death-resistant mutants, we searched for clones that survive heat-ramp and grow on agar plating (as opposed to USA300 which no longer show colony growth from replica plating) and a live/dead ratio that is higher than our death-resistant mutant, *cidA* control. After the heat-ramp exposure, we identified 74 mutants that were death-resistant. While we cannot pinpoint a specific gene to be the ultimate regulator of cell death, we generated a list of potential regulators of cell death. In order to identify core genes and pathways involved in cell death, we exposed the transposon mutant library to acetic acid stress.

Acetic acid has been shown to induce cell death in *S. aureus* and in yeast as well [15, 16, 19, 23]. Upon treatment with acetic acid (6 mM) overnight, 27 (*hsdR1, hslO, nsaS, sspA, folD, mfd, vraF, kdpB*, USA300HOU_2684, 0868, 0369, 0420, 1154, 0142, 0930, 2590, 0997, 2559, 0044, 2004, 1209, 0152, 2455, 0154, 2386, 0232, 0350) out of the 74 heat-ramp resistant mutants were also acetic acid resistant (Fig. 1). A majority of the genes identified encode transporters (n=9), involved in transcription (n=4), metabolism (n=3), peptidases (n=2) and phosphatases & kinases (n=2). For genes encoding transferases and proteins involved in stress response, nucleic acid synthesis and protein synthesis, one candidate was found in each category.

**Figure 1.**
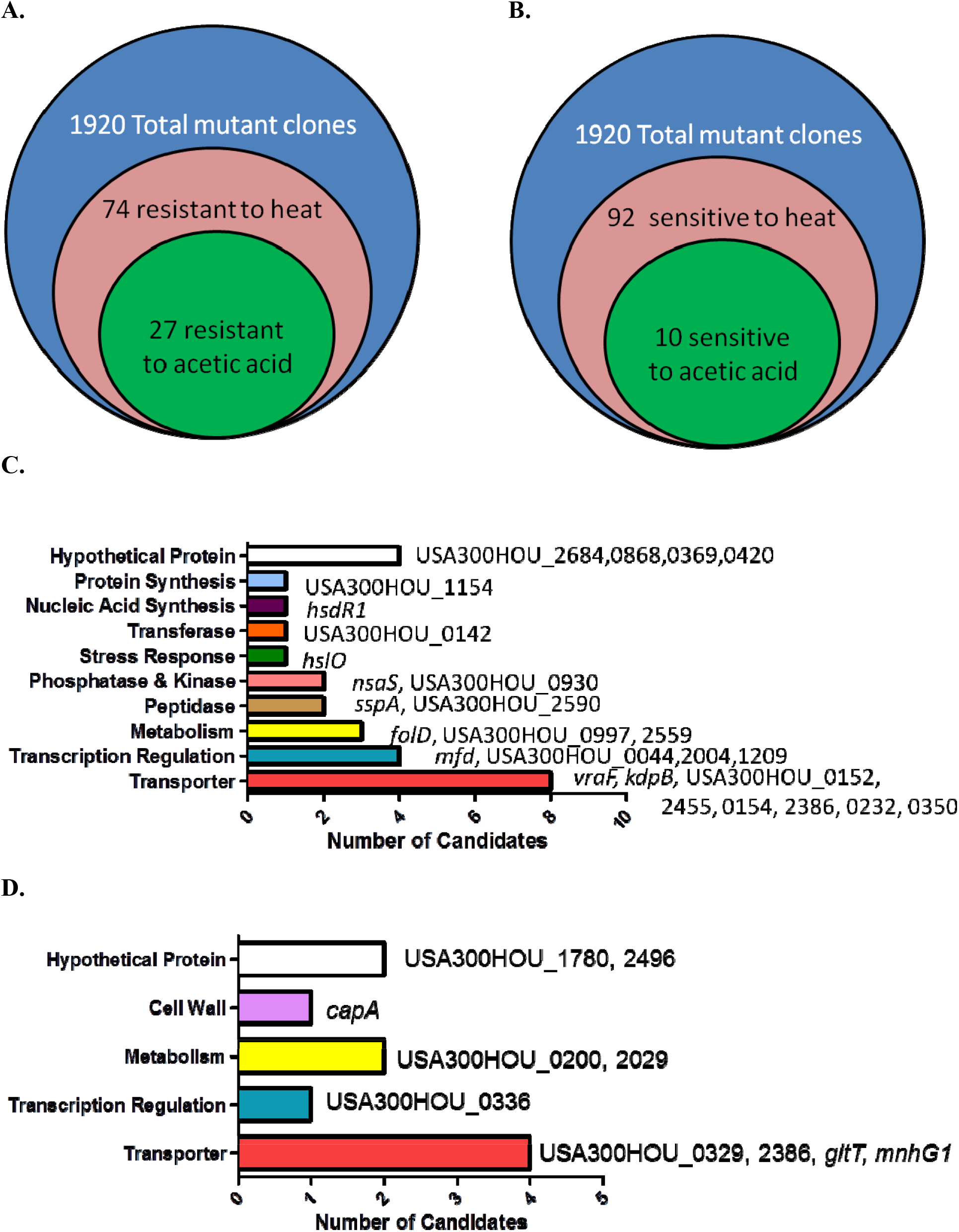
Identification of genes involved in causing bacterial cell death-resistance and cell death-sensitivity. (A) Summary of the number of candidates identified from the Nebraska Transposon Mutant Library (NTML) as being more death-resistant (A) and death-sensitive (B) in heat-ramp and acetic acid stress. The breakdown of the respective categories of genes whose mutations cause death-resistance (C) and death-sensitivity (D) to both stresses.

### Death-resistant strains are more virulent in vivo

Upon infection inside a host, in addition to drug stress, the bacteria are exposed to various types of stresses such as oxidative stress especially in the phagosomes of immune cells. We then tested if our death-resistant mutants were more virulent in causing an infection inside the host. After infection of *C. elegans* with the top 4 death-resistant mutants (*folD*, USA300HOU_0997, sspA, USA300HOU_0232) and parental strain USA300, we observed that all four mutants significantly decreased the survival of the *C. elegans* and killed the worms faster than USA300 (Fig. 2). By 2 days post-infection, the survival of worms infected with our death-resistant mutants had a survival rate of 36% or lower while worms infected with USA300 had a survival of 50%. The most virulent strain was Tn::USA300HOU_0232, a mutation in an iron transporter, as it caused the greatest mortality in the worms, resulting with only 22% survival of the worms by day 2. Our data suggest that bacteria that are more death-resistant could potentially cause more serious infections.

**Figure 2.**
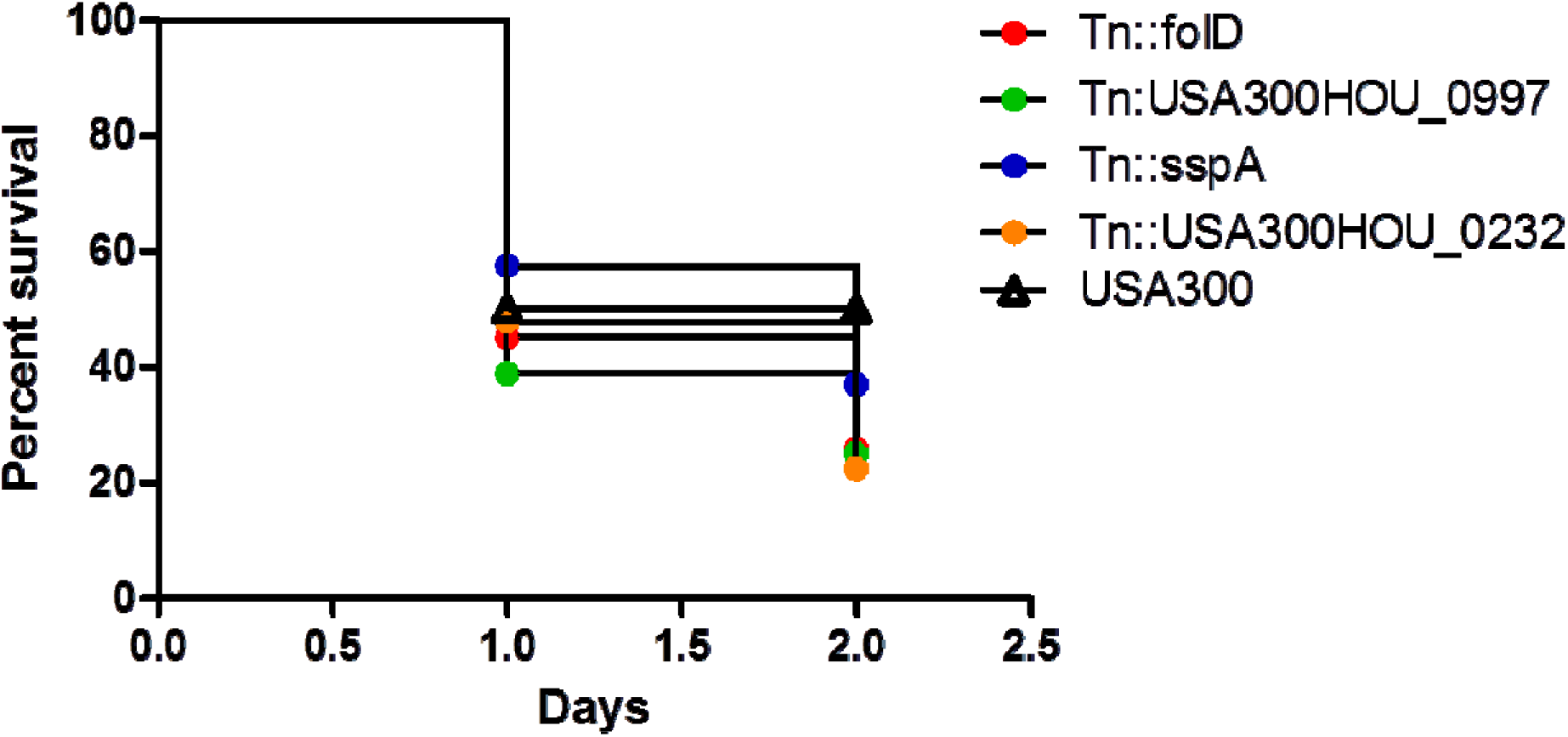
Strains that are more death-resistant caused more virulent infections in *C. elegans*. *C. elegans* (n=20-30) infected (10^6^ CFU) with mutants showing resistance to cell death showed increased mortality than *C. elegans* infected with parental strain USA300.

### Identification of genes important for cell death sensitivity revealed the importance of glutamate biosynthesis in cell death

In order to fully understand the regulatory networks of cell death pathways, it is crucial to examine the genes whose mutations cause the cells to be more sensitive to stress as well. Using the data from our two screens (heat-ramp and acetic acid stress), we adjusted the parameters of data analysis to distinguish mutants that were cell death sensitive. Unlike the cut-offs mentioned previously, cell death sensitive mutants were identified as having no viable growth on agar media and a live/dead ratio that was lower than USA300.

Our screen revealed that 92 mutants were hypersensitive to the heat-ramp, of which 10 (*capA, gltT, mnhG1*, USA300HOU_1780, 2496, 0200, 2029, 0336, 0329, 2386) were also sensitive to acetic acid stress (Fig. 1). Transporters were the more abundant (n=4) followed by genes involved in metabolism (n=2), and lastly with genes involved in transcription (n=1) and cell wall synthesis (n=1). Two of the candidates were hypothetical proteins. Interestingly, two (USA300HOU_2029, *gltT*) of the top four death-sensitive mutants (USA300HOU_0329, USA300HOU_2029, USA300HOU_2386, *gltT*) harbored mutations pertaining to glutamate metabolism.

### Death-sensitive strains are less persistent in vitro; strains with mutations involved in glutamate metabolism are the least persistent

One of the reasons why *S. aureus* can cause persistent and recalcitrant infections is due to its ability to form persister cells. Persisters are dormant cells that are formed during stressed conditions and upon stress removal, the bacteria can revert back to a growing state and consequently, cause a relapse in infection [24]. Bacterial persistence can also be viewed as cells with a strong anti-death program. Given now that we have identified genes whose mutation renders the bacteria death-sensitive, we then wanted to know if these mutations also lead to defective in persistence, forming lower amounts of persister cells. Such mutations could then potentially be drug targets for clearing persistent infections.

Persisters are induced by treating stationary phase bacteria with high concentrations of bactericidal antibiotics (usually at least 10 X MIC). Cells are then washed to rid of stress and plated on solid medium with no drug for CFU enumeration [25]. We exposed the top 4 death-sensitive mutants to bactericidal antibiotics with different mechanism of actions: gentamicin, meropenem, rifampin, and moxifloxacin. Upon 6-days post exposure of antibiotics, all 4 mutants showed a defect in persistence when exposed to all different classes of antibiotics (Fig. 3). The amount of persisters is dependent on the type of stress which can be seen here since the absolute amount of persisters changes among the drugs tested [24]. However, the overall amount of persister cells formed by the death-sensitive mutants was significantly lower than USA300. The defect in persistence was the most prominent for gentamicin stress. Under gentamicin stress, mutations in *gltT* and USA300HOU_2029, both involved in glutamate metabolism, were completely killed by 4 day-post exposure while USA300 still had over 10^7^ CFU/ml. In non-stressed conditions, no growth defects and decrease in viability of cells were observed.

**Figure 3.**
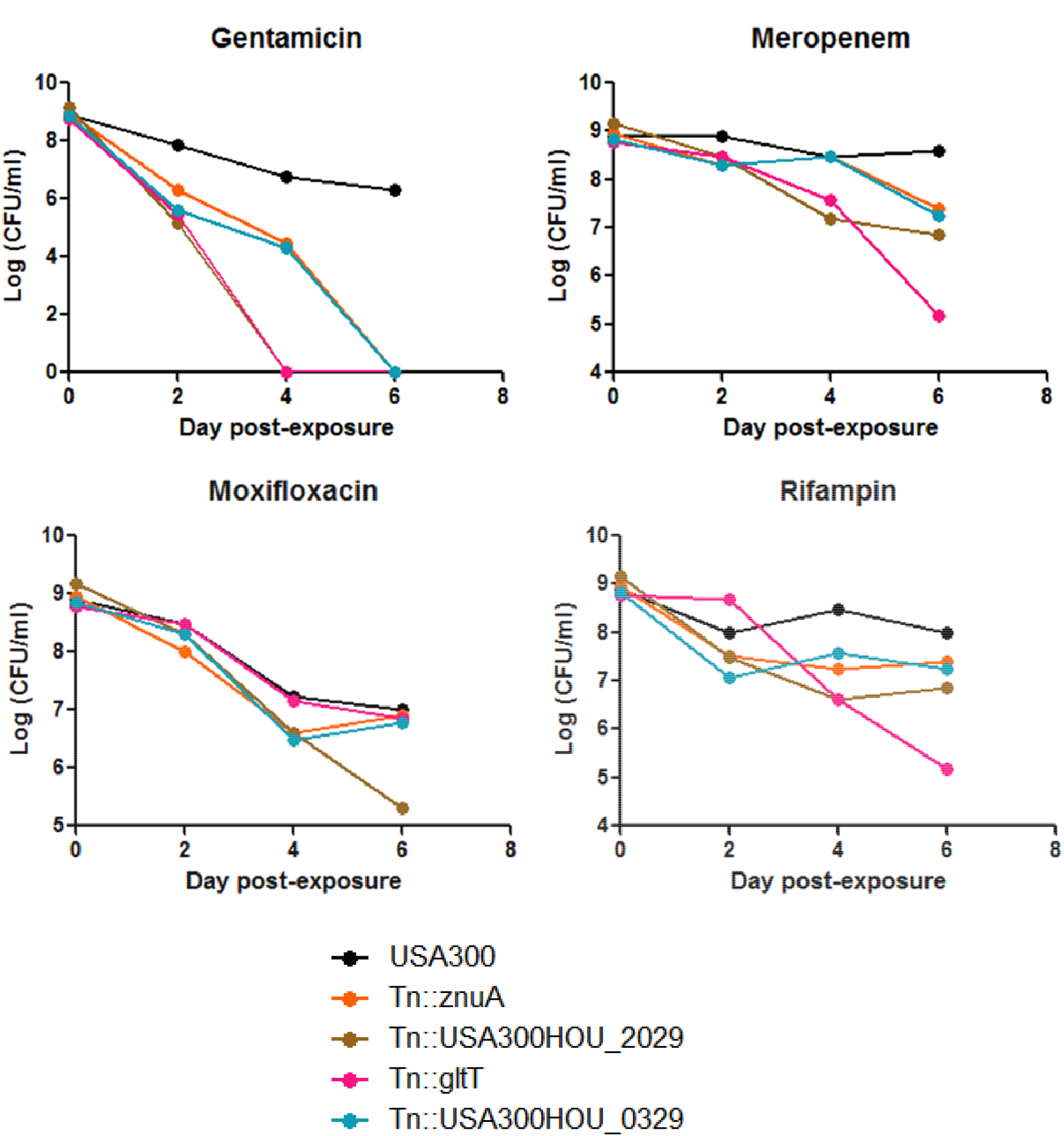
Strains that are more death-sensitive also show decreased persistence to bactericidal antibiotics. Mutants that were more cell-death sensitive showed defective persistence to gentamicin (60 μg/ml), meropenem (20 μg/ml), moxifloxacin (40 μg/ml), and rifampin (2 μg/ml) upon prolonged drug exposure up to 6 days.

## Discussion

To our knowledge, this is the first comprehensive study to identify genes and pathways that play a role in anti-death and pro-death programs in *S. aureus*. These data suggest that not only are transporters important in regulating cell death pathways in *S. aureus*, but in particular, glutamate metabolism and glutamate transport are important for transformation of a bacteria cell into a both a more death-resistant and a persistent phenotype under stressed conditions. Our findings show that mutations in genes involved in intracellular glutamate level regulation such as USA300HOU_2029 and *gltT* can decrease cell viability and persistence under antibiotic stresses, and environmental stresses such as temperature changes (heat-ramp) and low pH (acetic acid).

While other studies have searched for cell death genes in *S. aureus* [14, 15, 26, 27], the specific pathway of glutamate metabolism has never been identified as core cell death proteins. Although further research is needed to explore how glutamate metabolism plays a protective role in cell death, hypotheses based on what is currently known about cell death can offer insights into how intracellular glutamate levels could fit in the program of *S. aureus* cell death. Sadykov et al. identified that inactivation of the phosphotransacetylase-acetate kinase (Pta-AckA) pathway which normally generates acetate from acetyl-CoA leads to cell death in *S. aureus* under glucose and oxygen excess [26]. In bacteria, glutamate fermentation can occur via 3-methylaspartate which produces pyruvate followed by acetyl-CoA. Considering that acetate can be produced from acetyl-CoA, our findings may help explain the events that occur upstream of Pta-AckA activation [28]. Additionally, the lethality induced by cidal antibiotics has been shown to be due to ROS generation and radical generation from the Fenton reaction suggesting that death mechanisms result in oxidative responses within the cell [5]. In *Francisella*, glutamate transporter GadC has been shown to neutralize reactive oxygen species [29]. A study performed to evaluate the bactericidal effect of CO-releasing molecules (CO-RMs) showed that CO-RMs stimulated the production of intercellular ROS in the bacteria which was abolished when glutamate was supplemented to the culture [30]. Thus, increased glutamate levels in the stressed cells may protect the cell from ROS-mediated cell death under stress.

Findings from our previous screens performed to identify genes involved in persistence to rifampicin [25] showed that genes *gltS, gltD, gltA*, all of which are involved in glutamate synthesis, were important. Intriguingly, the protein ArgJ was shown to be a potential core regulator for *S. aureus* persistence in various stresses (different classes of drugs, heat, and low pH) [31]. Glutamate is the substrate for ArgJ (Fig. 4) and since mutants with impaired glutamate biosynthesis and transport showed both death-sensitive and defective persistence phenotypes, it can be speculated that glutamate can be the main driver of anti-death or equivalent to elevated persistence programs where arginine synthesis is a potential downstream effector pathway. Further biochemical studies and metabolomic studies are needed to understand how glutamate and arginine are involved in cell death.

**Figure 4.**
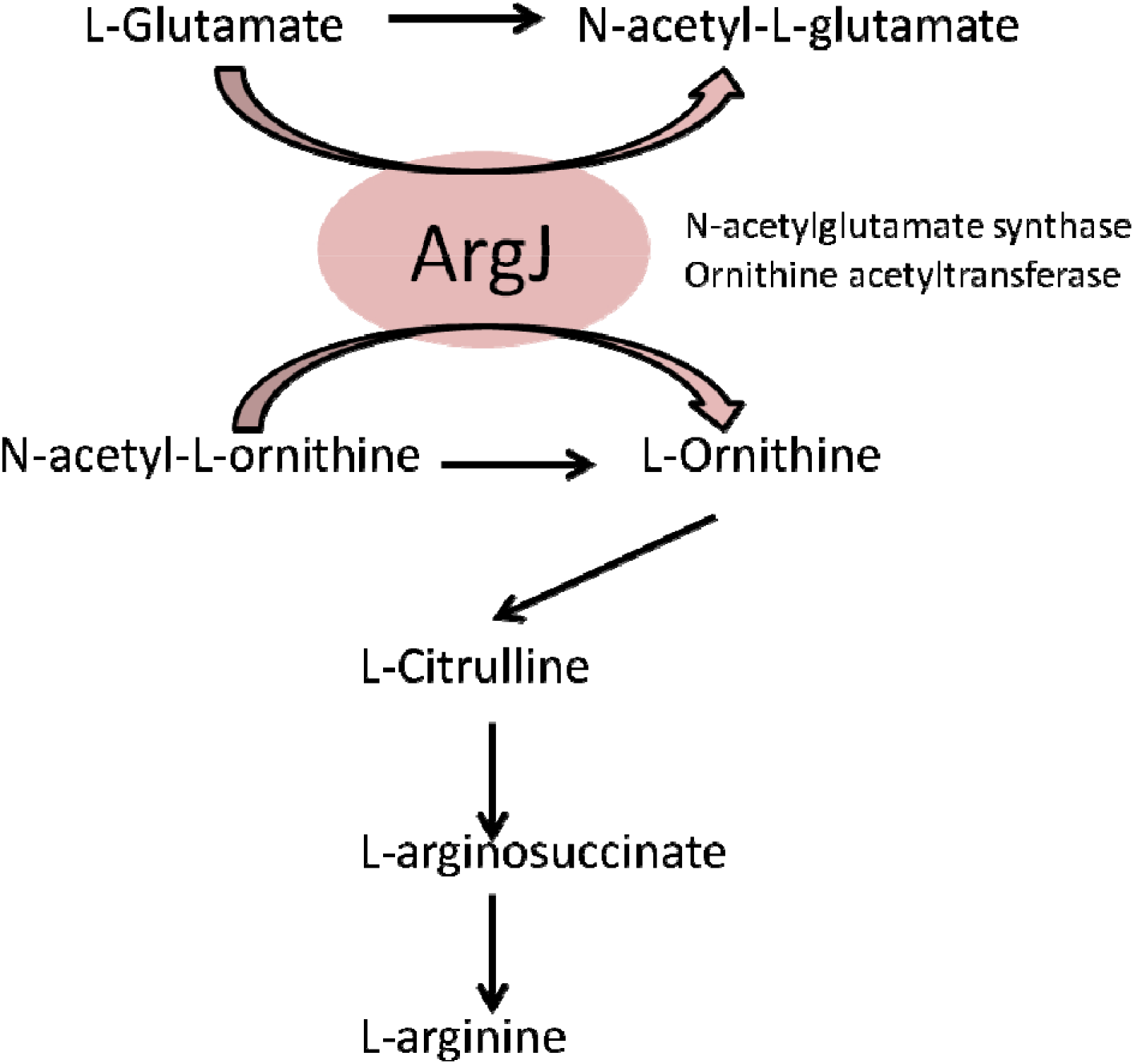
Proposed model of cell-death pathway through glutamate and arginine metabolism

Our screen also identified mutants whose phenotypes are cell death resistant and when the top four death-resistant mutants were tested in an *in vivo C. elegans* model, we found that they are indeed more virulent than the parental strain USA300 (Fig. 2). The mutant that was the most virulent harbored a mutation in USA300HOU_0232 which encodes an iron transporter. Similarly, the *nsaS* mutant was also identified to be death-resistant from our screens and studies in *S. aureus* have shown that NsaS is part of the cell-envelope two-component sensing system[32, 33]. In a mutant of *nsaS* in *S. aureus*, there is decreased association with metal ions on the cell surface and as a result, the intracellular concentration of metal ions was reduced [32]. Considering how ROS may play a role in cell death [5], a mutated iron transporter may cause the cells to be death-resistant by decreasing the amount of ROS inside the cell [34] as the transporter limits the amount of iron needed for the Fenton reaction which can damage DNA and cause cell death. USA300HOU_0997 encodes an autolysin family protein that plays a role in peptidoglycan turnover [35]. Gene expression of autolysins have been implicated in increasing cell wall stability under stressed conditions [36]. In vancomycin resistant strains, autolysin genes were downregulated which led to changes in membrane charges and thickness of the cell wall enabling the cell to survive treatment of cell wall inhibitors [37]. It can be speculated here that decreased cell division and peptidoglycan turnover in our USA300HOU_0997 mutant may lead to death resistance. FolD is a bifunctional protein that allows the production of continuous tetrahydrofolate which is a key metabolite for amino acid and nucleic acid biosynthesis [38]. The catalytic step of FolD is a checkpoint that regulates folate production [39]. As such, a mutant of *folD* may disturb the negative feedback of folate synthesis and continuously produce important metabolites and nucleotides for continued survival under stressed conditions. For example, MthfR is a Methylenetetrahydrofolate reductase in the same pathway as FolD and has been shown to be important for growth in *S. aureus* [40]. Lastly, the SspA protein encodes for the V8 serine protease that cleaves fibronectin binding proteins. Serine proteases such as SspA are under the *agr* quorum-sensing control operon [41] which coordinates a series of gene expression cascades to withstand stresses such as oxidative stress and the human immune response and facilitate bacterial cell growth and pathogenesis [42, 43]. It has been shown that *agr* negative cells do not undergo cell death as rapidly and were more resistant to cell lysis due to the increased expression of RNAIII [44, 45]. The increased mortality of *C. elegans* infected with the SspA mutant suggests that downregulation of *agr* gene could potentially be involved in causing cell death resistance because *agr*-negative *S. aureus* strains can be found in hosts with chronic infections and cause infections with increased mortality compared to those infected with *agr*-positive strains [46].

Interestingly, our screen revealed that mutants of *hslO* and *mfd* were more death-resistant which may appear contradictory given that expression of HslO and Mfd are protective for the cell during stress [47, 48]. HslO (or Hsp33) has been shown to be upregulated especially during oxidative stress and that HslO levels were decreased when *S. aureus* cells were transitioning to slow/non-growing status [49]. It is well known that Mfd is involved in prevent DNA damage from oxidative stress, immune response, and drugs [47, 50]. Decreased expression of Mfd in *S. aureus* led to decreased biofilm formation [51] and a *mfd* mutant of *C. difficle* has increased toxin production [52]. For *B. subtilis*, Mfd-deficient cells formed less endospores [53] and resulted in a 35-fold overexpression of OhrR, a transcription factor involved in peroxide stress response [54]. Given that Mfd has many nonconical roles, we speculate that under stressed conditions decrease of expression in our *mfd* mutant may allow expression of another transcription factor (such as the OhrR) to activate stress response pathways. It is also possible that the contradictory phenotypes of the *hslO* and *mfd* mutants are due to compensatory mutations in the *hslO* and *mfd* mutants, which are known to occur in loss of function mutants in yeast [55].

As with any mutant library, secondary mutations may have occurred in some mutants that could affect the phenotypic outcomes. For example, our speculation as to why a *hsdR* mutant is more death-resistant could also be associated to the genetic background of the mutant. Inactivation of HsdR, an endonuclease of the type 1 restriction system in *S. aureus*, allowed *S. aureus* to become readily transformable due to decreased cleavage of foreign DNA [56, 57]. In our *hsdR* mutant, it could be possible that secondary mutations or genetic modifications have occurred in the genome that caused the death-resistant phenotype. While we performed whole genome sequencing on our mutant clones to confirm the location of the transposon, the possibility of secondary mutations playing a role in the phenotypes observed here have yet to be explored. However, we performed the screen twice followed by individual confirmation. The same subculture from the same stock was also used each time to ensure reproducibility among all replicates. Meanwhile, even if there is a secondary mutation, the decreased expression of the specific genes identified here due to the transposon insertion would still be a valid part of the mechanisms of cell death.

Our work adapted assays that are used in the yeast community to identify genes involved in yeast cell death [16–19]. In the high-throughput screen consisting of yeast mutants and even in yeast strains of different backgrounds, categories of genes that were redundant among the studies include carbohydrate metabolism, transcription factors, and ion transport [18, 58, 59]. Interestingly, amino acid transport was the most significantly enriched term for genes involved in positive regulation of acetic acid-induced death; two of the identified transporters were involved in metabolism of glutamate (GDH1 and GDH2) [18]. A similar approach of a “heat-ramp” stress was used in one study on *B. subtilis* using a water bath [60]. Their study revealed that heat shock proteins, sporulation, competence, and carbon metabolism were important. While we did not identify similar genes, carbon metabolism was identified in both our screens. Heat shock proteins were heavily enriched in their study but not ours which can be attributed to the candidates in our library which only contain mutants of non-essential genes and only 2 candidates out of 1,952 were heat shock proteins [61].

Despite the significant findings of this study, there are some limitations. First, while we generated a list of the mutants that are death-sensitive and death-resistant to both heat ramp and acetic acid stress, the screens performed here are only a snapshot of what is happening in the cell. The phenotypes seen here were determined by the condition of our assays and can be affected by the level of stress (e.g. concentration of the acetic acid, ramping time and temperature for the heat ramp) [17]. Although we included both a wild type USA300 strain and a CidA mutant, a known death-resistant mutant [15, 62], to optimize our conditions, it is important to recognize that our screen may not be comprehensive in finding all the mediators of cell death. While our goal was more conservative and was to search for core regulators, the significant role of other genes and pathways involved in cell death that were not classified as core regulators should not be undermined. Our screen only explored cell death at the DNA level and further protein studies would provide more comprehensive insight into the effects of gene transposon insertions in regulating cell death. Even though this mutant library contained all the non-essential genes in USA300, we cannot overrule the fact that cell death is an important program for any living organism and cell death regulators may be essential genes which are not included in our library.

Given that we know the killing activity of antibiotics extend beyond the drug-target interactions [5], understanding the effector proteins and downstream events of bacterial cell death can help provide novel drug treatment approaches for bacterial infections. For example, cell death can be artificially induced by disruption of the negative regulator of cell death in a similar fashion to how apoptotic pathways are exploited to treat cancer cells. Recently, it was shown that extracellular death peptides from both *E. coli* and *P. aeruginosa* can induce toxic endoribonucleolytic activities of MazEF in *M. tuberculosis* suggesting promising therapeutic outcomes upon manipulation of the cell death mediators in bacteria [63]. Similarly, since ROS formation serves an important role in the lethality of cidal antibiotics, a drug combination with metabolic perturbations may enhance killing of currently used antibiotics. For example, causing defects in peroxide-detoxifying enzymes have been shown to increase antimicrobial lethality [64]. On the other hand, it is of concern that more than half of the population in the US consumes nutritional supplements such as vitamins which are antioxidants [65] and the potential antagonistic effects of an antioxidant diet in a patient taking antibiotics will require further investigation.

In conclusion, we report the molecular basis of cell death in *S. aureus*. To our knowledge, this is the first report on identification of *S. aureus* cell death genes from a whole genome perspective. Our extensive screen also offers insights to common core mechanisms that are relevant to not only cell death but bacterial persistence, a phenomenon that’s at the core of recalcitrant infections and biofilm formations. Our studies provide insights to possible new drugable targets, biomarkers for recalcitrant infections for diagnostic purposes and novel vaccine targets for prevention of bacterial infections. The similarity in functional groups found between our study and other yeast studies suggest that our work also sheds light into cell death pathways of eukaryotic systems such as pathogenic fungi and cancer stem cells.

## Methods

### Culture Media, Antibiotics, and Chemicals

Meropenem, moxifloxacin, rifampicin, gentamicin, and erythromycin were obtained from Sigma-Aldrich Co. (St. Louis, MO, USA). Stock solutions were prepared in the laboratory, filter-sterilized and used at indicated concentrations. Bacterial strains used in this study include USA300 and the Nebraska-Transposon Mutant Library (NTML) [20]. *S. aureus* strains were cultured in tryptic soy broth (TSB) and tryptic soy agar (TSA) with the appropriate antibiotics and growth conditions. Transposon-insertion mutants grew in erythromycin (50 μg/ml), the antibiotic selective marker.

### Genetic Screen to identify cell death mutants

For the heat-ramp, we performed the assay as described [17]. Briefly, we normalized the concentration of the bacteria to OD600= 0.5 using PBS as the diluent. Then, we placed the samples in the thermocycler with a protocol following: 30°C for 1 minute, ramp from 30 °C to 62 °C with a step-interval of 0.5 °C per 30 °C seconds. For acid stress, acetic acid (6 mM) were added into stationary phase cultures and incubated overnight. To enumerate for cell counts, the mutant library was replica transferred to TSA plates to score for mutants that failed to grow after stress. For viability staining, SYBR Green I/PI staining was performed as described [21, 22]. SYBR Green I (10,000× stock, Invitrogen) was mixed with PI (20□mM, Sigma) in distilled H_2_O with a ratio of 1:3 (SYBR Green I to PI) in 100 ul distilled H2O and stained for 30 minutes in room temperature. Prior to the heat-ramp, the SYBR Green I/PI dye was added to the bacteria as the heat-ramp impaired the uptake of the dye. For the acetic acid stress, the dyes were added at the end of the exposure [17]. The green and red fluorescence intensity was detected using a Synergy H1 microplate reader by BioTek Instruments (VT, USA) at excitation wavelength of 485□nm and 538□nm and 612□nm for green and red emission, respectively. The live/dead ratio was calculated by dividing the green/red fluorescence.

### Persister Assays

Selected drugs were added to overnight stationary phase cultures for 6 days. At the selected time points, bacterial cultures were washed with 1 X PBS to remove stress, serially diluted, and plated onto TSA with no drugs for cell enumeration [25].

### Nematode-killing Assay

*C. elegans* N2 Bristol worms (Caenorhabditis Genetics Center) were synchronized to the same growth stage by treatment with alkaline hypochlorite solution as described [66]. Worms of adult stage were washed and suspended in bleaching solution with 5% hypochlorite for 9 minutes to lyse all the adult stages but keeping the eggs intact. Bleach was removed by centrifugation at 1,500 rpm for 1 minute and washed three times with M9 buffer. The pellet was incubated in M9 buffer at 20 °C with gentle agitation and proper aeration. L4 stage adult worms were obtained after 48 hours at 20 °C. For each assay, 20-30 worms were added to liquid M9 buffer supplemented with 5-Fluoro-2’-deoxyuridine (10 μM) to inhibit progeny formation. *S. aureus* (10^6^ CFU) that were grown overnight at 30 °C in TSB containing the appropriate antibiotics as needed were added into the buffer containing the worms. The samples were scored for live and dead worms every 24 hours.

**Table 1.** Top 4 genes whose mutations result in resistance to cell death in both heat-ramp and acetic acid stress

| Accession Number | Gene Name (if applicable) | Function | KEGG Ontology |
| --- | --- | --- | --- |
| USA300HOU_1008 | *folD* | Methylenetetrahydrofolate dehydrogenase; methenyltetrahydrofolate cyclohydrolase | Metabolism of cofactors and vitamins; One carbon pool by folate |
| USA300HOU_0997 |  | Bifunctional N-acetylmuramoyl-L-alanine amidase/mannosyl-glycoprotein endo-beta-N-acetylglucosaminidase | Metabolism |
| USA300HOU_0996 | *sspA* | Glutamyl endopeptidase | Cellular community prokaryotes; Quorum sensing; Serine Peptidases of chymotrypsin family |
| USA300HOU_0232 |  | Iron ABC transporter membrane binding protein | Environmental Information Processing; Membrane transport; ABC transporters; Mineral and organic ion transporters; Iron transporter |

**Table 2.** Top 4 genes whose mutations result in sensitivity to cell death in both heat-ramp and acetic acid stress

| Accession Number | Gene name (if applicable) | Function | KEGG Ontology |
| --- | --- | --- | --- |
| USA300HOU_0329 |  | ABC transporter-ATP binding protein | Protein families: Signaling and cellular processes; Transporters |
| USA300HOU_2029 |  | Amidohydrolase | Metabolism: Amino acid metabolism; Alanine, aspartate and glutamate metabolism |
| USA300HOU_2386 | *znuA* | Zinc transport system substrate-binding protein | Environmental Information Processing; Membrane transport; Metallic cation, iron-siderophore and vitamin B12 transporters; Zinc transporter |
| USA300HOU_2366 | *gltT, gltP* | Proton glutamate symport protein | Protein families: Signaling and cellular processes; Electrochemical potential-driven transporters |

## Acknowledgements

We gratefully acknowledge BEI Resources for provision of the NTML mutant library. We thank Zachary Stolp and Marie Hardwick for helpful discussions regarding the heat-ramp experiment. RY was supported by NIH training grant T32 AI007417.

---

**Supplementary Table 1.**
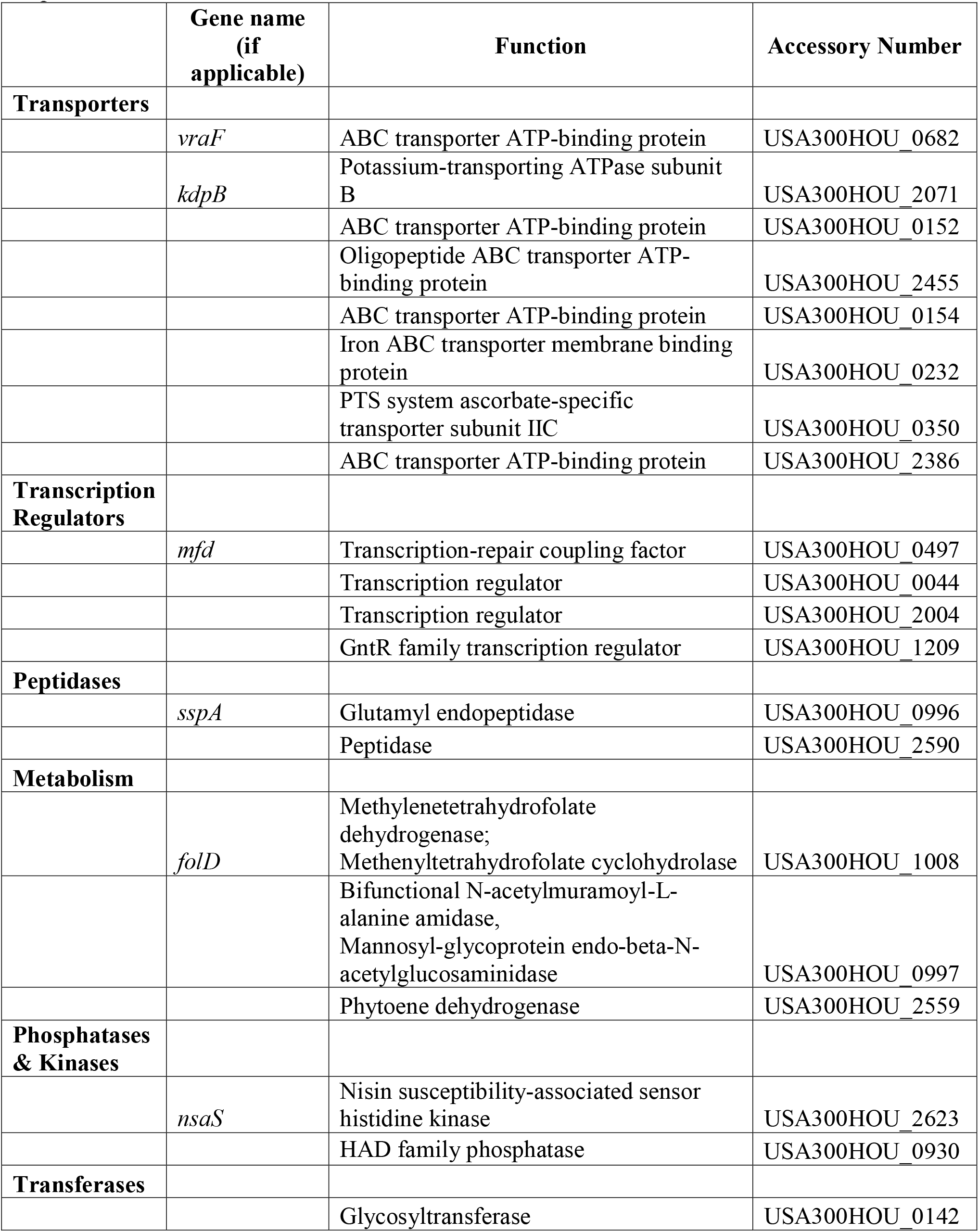
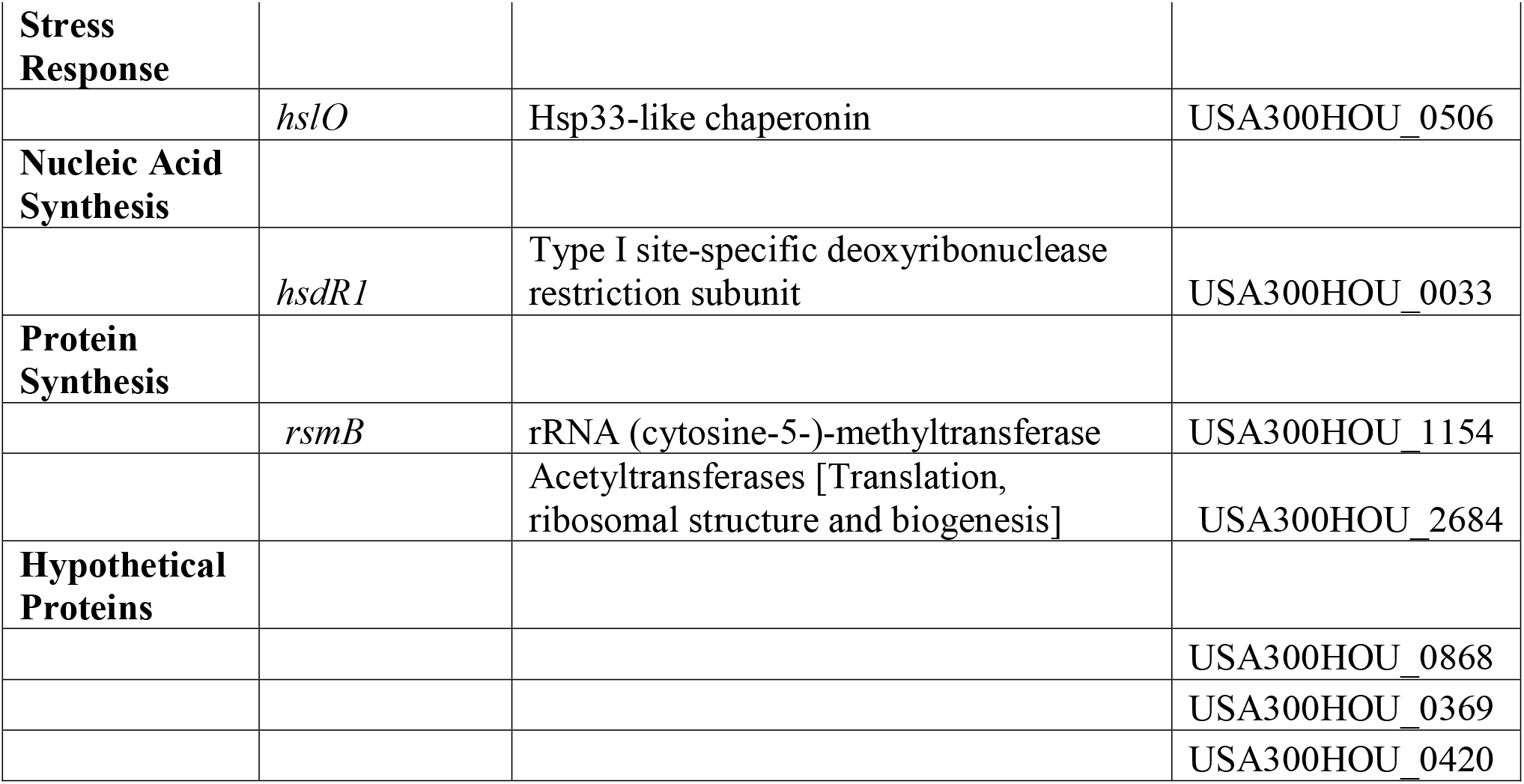
Genes whose mutations resulted in cell death resistance to both heat-ramp stress and acetic acid stress

**Supplementary Table 2.** Genes whose mutations resulted in more cell death after heat-ramp and acetic acid stress

|  | Gene name<br>(if applicable) | Function | Accessory Number |
| --- | --- | --- | --- |
| **Transporters** |  |  |  |
|  | *gltT* | Proton glutamate symport protein | USA300HOU_2366 |
|  | *mnhG1* | Monovalent cation antiporter subunit G | USA300HOU_0649 |
|  |  | ABC transporter ATP-binding protein | USA300HOU_0329 |
|  |  | ABC transporter ATP-binding protein | USA300HOU_2386 |
| **Metabolism** |  |  |  |
|  |  | Isochorismatase | USA300HOU_0200 |
|  |  | Amidohydrolase | USA300HOU_2029 |
| **Transcription** |  |  |  |
|  |  | Transcription regulator | USA300HOU_0336 |
| **Cell wall** |  |  |  |
|  | *capA* | Capsular polysaccharide biosynthesis protein | USA300HOU_2664 |
| **Hypothetical Proteins** |  |  |  |
|  |  |  | USA300HOU_1780 |
|  |  |  | USA300HOU_2496 |

